# Improved DNA-versus-Protein Homology Search for Protein Fossils

**DOI:** 10.1101/2021.01.25.428050

**Authors:** Yin Yao, Martin C. Frith

**Affiliations:** Graduate School of Frontier Sciences, University of Tokyo; Artificial Intelligence Research Center, AIST; Computational Bio Big-Data Open Innovation Laboratory (CBBD-OIL), AIST

## Abstract

Protein fossils, i.e. noncoding DNA descended from coding DNA, arise frequently from transposable elements (TEs), decayed genes, and viral integrations. They can reveal, and mislead about, evolutionary history and relationships. They have been detected by comparing DNA to protein sequences, but current methods are not optimized for this task. We describe a powerful DNA-protein homology search method. We use a 64×21 substitution matrix, which is fitted to sequence data, automatically learning the genetic code. We detect subtly homologous regions by considering alternative possible alignments between them, and calculate significance (probability of occurring by chance between random sequences). Our method detects TE protein fossils much more sensitively than `blastx`, and > 10× faster. Of the ~7 major categories of eukaryotic TE, three have not been found in mammals: we find two of them in the human genome, polinton and DIRS/Ngaro. This method increases our power to find ancient fossils, and perhaps to detect non-standard genetic codes. The alternative-alignments and significance paradigm is not specific to DNA-protein comparison, and could benefit homology search generally.

## 1 Introduction

Genomes are littered with protein fossils, old and young. They can be found by comparing DNA to known proteins: new transposable element (TE) families have been discovered in this way [25]. An interesting class of protein fossils comes from ancient integrations of viral DNA into genomes, enabling the field of paleovirology [17]. The DNA sequences of protein fossils often have similarity to distantly-related genomes (e.g. mammal versus fish), simply because the parent gene evolved slowly, so it is important to know that they are protein fossils in order to understand this similarity [28]. DNA-protein homology search is also used to classify DNA reads from unknown microbes, including nanopore and PacBio reads with many sequencing errors [16]. DNA-protein comparison can be used to find frameshifts during evolution of functional proteins [26], and programmed ribosomal frameshifts [35]. A more specialized and complex kind of DNA-protein comparison, outside this study’s scope, considers introns and other gene features to identify genes.

DNA-protein homology search is a classical problem with many old solutions [23, 30, 12, 15, 39, 13, 11, 20, 4, 22, 34]. A notable one is “three-frame alignment” [39], which we believe is the simplest and fastest reasonable way to do frameshifting DNA-protein alignment. Nevertheless, we can significantly improve DNA-protein homology search in these aspects:

– Better parameters for the (dis)favorability of substitutions, deletions, insertions, and frameshifts. Most previous methods use standard parameters such as the BLOSUM62 substitution matrix, which is designed for functional proteins, and likely completely inappropriate for protein fossils. We optimize these parameters by fitting them to sequence data.
– Instead of a 20 × 20 substitution matrix, use a 64 × 21 matrix (64 codons × 20 amino acids plus STOP). This allows e.g. preferred alignment of asparagine (which is encoded by `aac` and `aat`) to `agc` than to `tca`, which both encode serine.
– Incorporate frameshifts into affine gaps. Because gaps are somewhat rare but often long, it is standard to disfavor opening a gap more than extending a gap. However, most previous methods favor frameshifts equally whether isolated or contiguous with a longer gap.
– Detect homologous regions based on not just one alignment between them, but on many possible alternative alignments. This is expected to detect subtle homology more powerfully [1, 6].
– Calculate significance, i.e. the probability of such a strong similarity occurring by chance between random sequences. To this day, for ordinary alignment, BLAST can only calculate significance for a few hardcoded sets of substitution and gap parameters. We can do it for any parameters, for similarities based on many alternative alignments.

We also aimed for maximum simplicity and speed, inspired by three-frame alignment.

## 2 Methods

### 2.1 Alignment elements

We define a DNA-protein alignment to consist of: matches (3 bases aligned to 1 amino acid), base insertions, and base deletions. To keep things simple, insertions are not allowed between bases aligned to one amino acid. A deletion of length not divisible by 3 leaves “dangling” bases (Fig. 1): for simplicity, we do not attempt to align these (equivalently, align them to the amino acid with score 0).

**Fig. 1.**
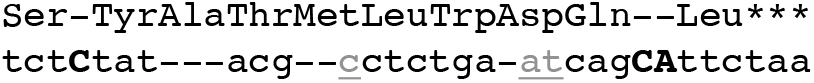
Example of a DNA-versus-protein alignment. *** indicates a protein end from translation of a stop codon. Insertions are bold uppercase. “Dangling” bases, left by deletions of length not divisible by 3, are underlined gray.

### 2.2 Scoring scheme

An alignment’s score is the sum of:

– Score for aligning amino acid *x* to base triplet *Y* : *S_xY_*
– Score for an insertion of *k* bases: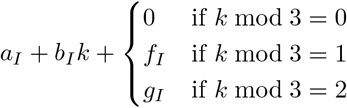
– Score for a deletion of *k* bases: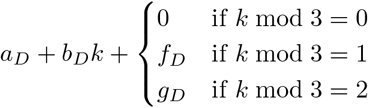

This scheme extravagantly uses 4 frameshift parameters (*f_I_*, *g_I_*, *f_D_*, *g_D_*), because it’s based on a probability model with 4 frameshift transitions (Fig. 2), and we can’t think of a good way to simplify the model. Overall, our alignment scheme is similar to FramePlus [13] and especially to aln [11].

**Fig. 2.**
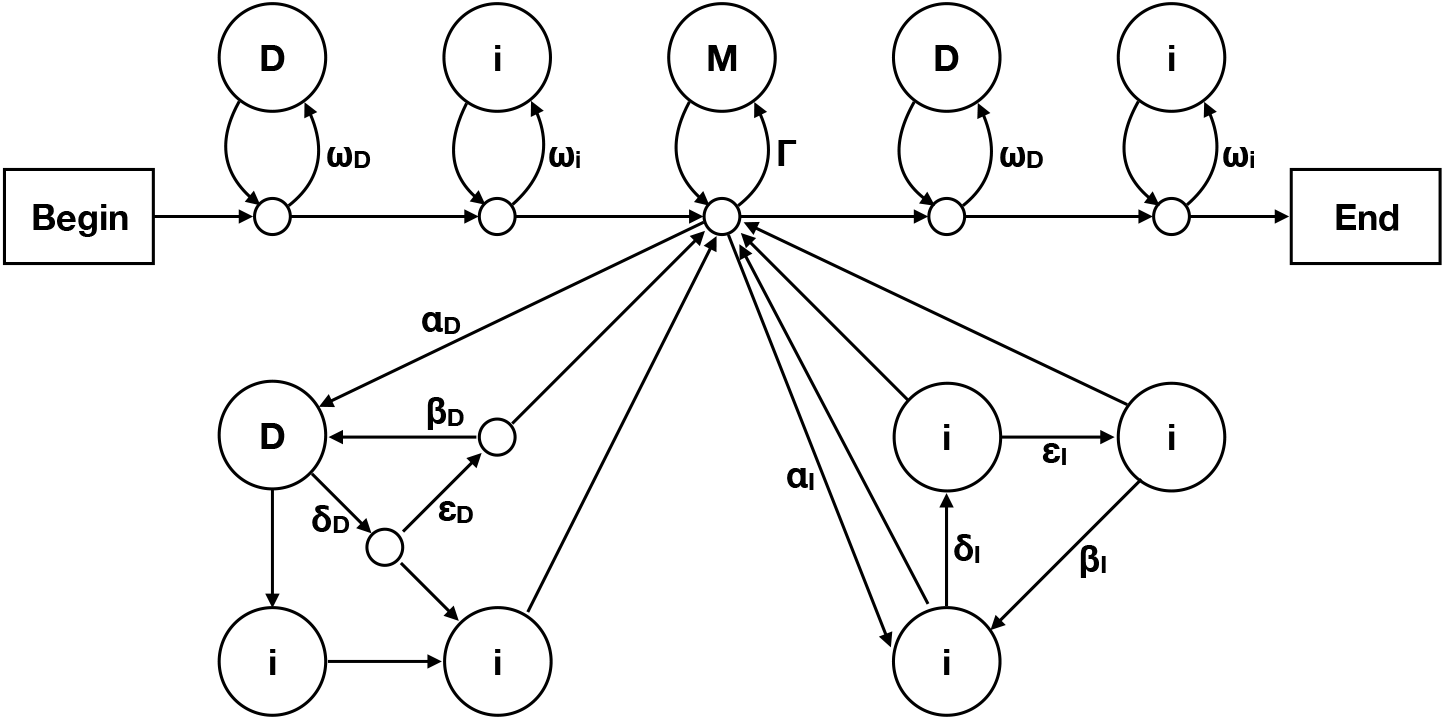
A probability model for related DNA and protein sequences. The arrows are labeled with probabilities of traversing them. Each pass through an **i** state generates one base *y* ∈ {`a, c, g, t`}, with probabilities *ψ_y_*. Each pass through a **D** state generates one amino acid *x*, with probabilities *ϕ_x_*. Each pass through the **M** state generates one amino acid *x* aligned to three bases *Y* = *y*_1_*y*_2_*y*_3_, with probabilities *π_xY_*. The two bottom-left **i** states correspond to “dangling” bases.

### 2.3 Finding a maximum-score local alignment

A basic approach is to find a maximum-score alignment between any parts of a protein sequence *R*_0_ … *R*_M−1_ and a DNA sequence *q*_0_ … *q*_*N*−1_. Let *Q_j_* mean the triplet *q_j_, q*_*j*+1_, *q*_*j*+2_. We can do these calculations for 0 ≤ *i* ≤ *M* and 0 ≤ *j* ≤ *N* :

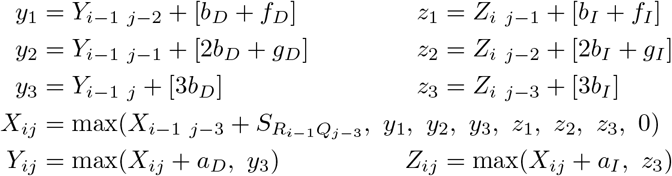

The boundary condition is: if *i* < 0 or *j* < 0, *X_ij_* = *Y_ij_* = *Z_ij_* = −∞. The maximum possible alignment score is max(*X_ij_*), and an alignment with this score can be found by a standard traceback [5].

For each (*i, j*) this algorithm retrieves 7 previous results, and performs 9 pairwise maximizations and 9 additions (which could be reduced to 6 additions if each insertion cost equals its corresponding deletion cost). This is slightly slower than three-frame alignment, which retrieves 5 previous results and performs 7 pairwise maximizations and 6 additions.

### 2.4 Probability model

The preceding algorithm is equivalent to finding a maximum-probability path generating the sequences, through a probability model (Fig. 2). The score and model parameters are related like this:

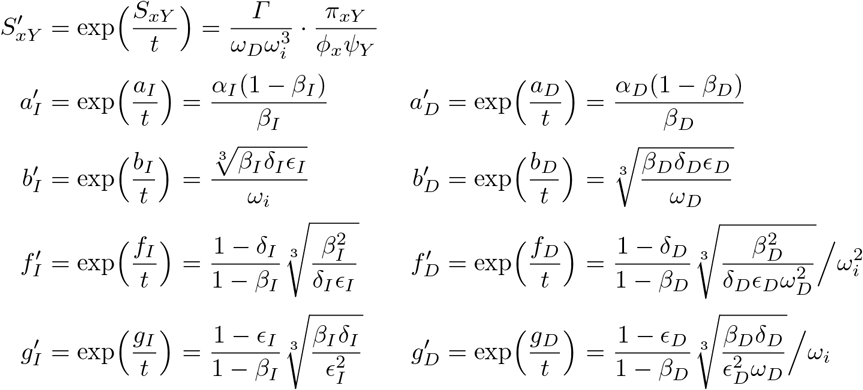

Here *ψ_Y_* is defined to be *ψ*_*y*1_ *ψ*_*y*2_ *ψ*_*y*3_, and *t* is an arbitrary positive constant (because multiplying all the score parameters by a constant makes no difference to alignment). An alignment score is then: *t* ln[prob(path & sequences)/ prob(null path & sequences)], where a “null path” is a path that never traverses the *Γ*, *α_D_*, or *α_I_* arrows [10].

#### Balanced length probability

A fundamental property of local alignment models is whether they are biased towards longer or shorter alignments [10]. If *ω_D_* and *ω_i_* are large (close to 1) and *Γ* + *α_D_* + *α_I_* is small, there is a bias in favor of shorter alignments. In the converse situation, there is a bias towards longer alignments. It can be shown (using the method of [10]) that our DNA-protein model is unbiased when

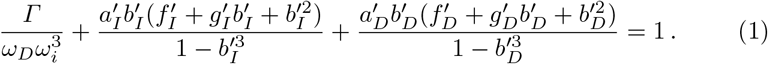

### 2.5 Sum over all alignments passing through (*i, j*)

To find subtly homologous regions, we should assess their homology without fixing an alignment [1, 6]. In other words, we should use a homology score like this: *t* ln [Σ_paths_ prob(path & sequences)/prob(null path & sequences)]. However, if the sum is taken over all possible paths, we learn nothing about location of the homologous regions, which is important if e.g. the DNA sequence is a chromosome. There is a kind of uncertainty principle here: the more we pin down the alignment, the less power we have to detect homology. As a compromise, we sum over all paths passing through one (protein, DNA) coordinate pair (*i, j*). This has two further benefits: it is approximated by the seed-and-extend search used for big sequence data, and we can calculate significance.

To calculate this sum over paths, we first run a Forward algorithm for 0 ≤ *i* ≤ *M* and 0 ≤ *j* ≤ *N* :

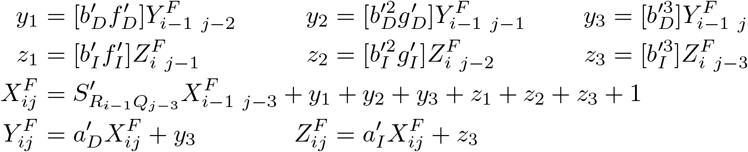

The boundary condition is: if *i* < 0 or *j* < 0, 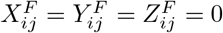. We then run a Backward algorithm for *M* ≥ *i* ≥ 0 and *N* ≥ *j* ≥ 0:

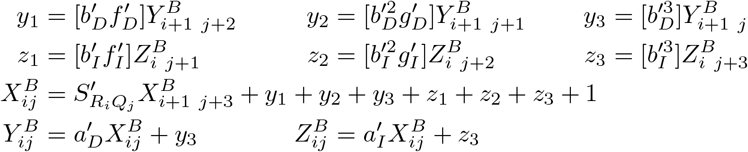

The boundary condition is: if *i* > *M* or *j* > *N*, 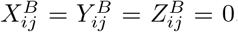. Finally, 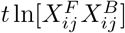 is the desired homology score, for all paths passing through (*i, j*).

### 2.6 Significance calculation

The just-described homology score is similar to that of “hybrid alignment”, which has a conjecture regarding significance [37]. (Hybrid alignment sums over paths ending at (*i, j*), instead of passing through (*i, j*).) We make a similar conjecture. Suppose we compare a random i.i.d. protein sequence of length *M* and letter probabilities *Φ_x_* to a random i.i.d. DNA sequence of length *N* and triplet probabilities *ψ_Y_*. We conjecture that the score 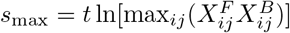 follows a Gumbel distribution:

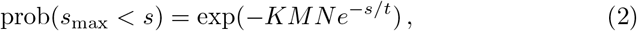

in the limit that *M* and *N* are large, provided that:

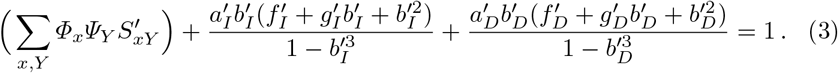

Equation 3 is analogous to Equation 27 or 28 in [37], see also [10]. In practice, we assume that *Φ_x_* = *ϕ_x_* and *ψ_Y_* = *ψ_Y_*, which makes Equation 3 equivalent to Equation 1.

This conjecture leaves one unknown Gumbel parameter *K*. We estimate it by brute-force simulation of 50 pseudorandom sequence pairs [38], with *Φ_x_* = *ϕ_x_*, *ψ_Y_* = *ψ_Y_*, *M* = 200 and *N* = 602, which takes zero human-perceptible run time.

### 2.7 Seed-and-extend heuristic

To find homologous regions in big sequence data, we use a BLAST-like seed-and-extend heuristic (Fig. 3) [2]. We first find “seeds”: we currently use exact-matches (via the genetic code), which can be sensitive if short, but we could likely get better sensitivity per run time with inexact seeds [27, 31]. Our seeds have variable length: starting from each DNA base, we get the shortest seed that occurs ≤ *m* times in the protein data [19]. We then try a gapless *X*-drop extension in both directions, and if the score achieves a threshold *d*, we try a “Forward” extension in both directions.

**Fig. 3.**
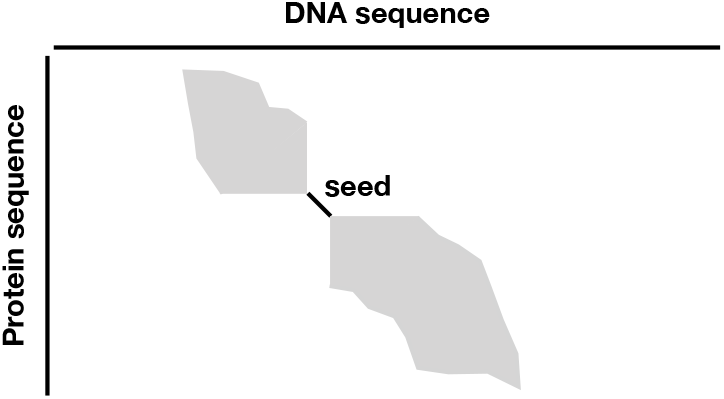
Sketch of seed-and-extend heuristic for homology search.

We use our Forward algorithm, modified for semi-global instead of local alignment. In each direction, we sum over alignments starting at the seed and ending anywhere: thus the algorithm’s +1 is done only at the first (*i, j*) next to the seed, and we accumulate the sum 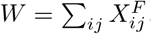. We run this algorithm in increasing order of antidiagonal (3*i* + *j*) on the seed’s right side (decreasing order on the left side). If 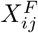 is less than a fraction *f* of *W* accumulated over previous antidiagonals, we stop extending, which defines the boundary of the gray region in Fig. 3. The final homology score is *t* ln[*W*_left_] + seed score + *t* ln[*W*_right_].

Sum-of-path algorithms are prone to numerical overflow [5]. To prevent that: once per 32 antidiagonals, we multiply all the *X^F^*, *Y^F^*, and *Z^F^* values in the last six antidiagonals by a scaling factor of 1/*W*.

A score with no alignment is disconcerting, so we get a representative alignment by a similar semi-global modification of our maximum-score alignment algorithm. To avoid redundancy, we prioritize homology predictions by score (breaking ties arbitrarily), and discard any prediction whose representative alignment shares an an (*i, j*) left or right end with a higher-priority prediction.

### 2.8 Fitting substitution & gap parameters to sequence data

We can fit the parameters to some related (unaligned) DNA and proteins, by an iterative Baum-Welch algorithm [5]. We implemented two versions of this: an exact *O*(*MN*) version, and a seed-and-extend version. The seed-and-extend version, at each iteration, finds significantly homologous regions (with -`K1` filtering, see below) and gets expected counts from the seeds and extend regions (gray areas in Fig. 3). It does not infer *ϕ_x_*, *ψ_y_*, *ω_D_*, or *ω_i_* in the usual way: at each iteration, it sets *ϕ_x_* = Σ_*Y*_ *π_xY_*, *ψ_y_* = Σ_*xij*_ (*π_x yij_* + *π_x iyj_* + *π_x ijy_*)/3, and 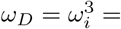 the value that satisfies Equation 1 (found by bisection with bounds 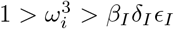 and 1 > *ω_D_* > *β_D_δ_D_є_D_*). We set *t* = 3/ ln[2] to get scores in third-bit units.

## 3 Results

### 3.1 Parameter fitting

We applied our *O*(*MN*) fitting to a set of human processed pseudogenes and their parent proteins from Pseudofam [21]. To avoid bias, we began the first iteration with *π_xY_* = 1/(21 · 64). The fitting discovered the genetic code: for each codon *Y*, its encoded amino acid has maximum *S_xY_*.

Sometimes, our fitting had an undesirable feature: the *S_xY_* values for some `cg`-containing codons were all negative. This is presumably due to the well-known depletion of `cg` in human DNA, which can be captured in *π_xY_* but not *ψ_y_*. As an ad hoc fix, we set *ψ_Y_* = Σ_*x*_ *π_xY_* (after *O*(*MN*) fitting, and at each iteration of seed-and-extend fitting).

Next, we applied our seed-and-extend fitting to human chromosome 21 (hg38 chr21) and transposable element (TE) proteins from RepeatMasker 4.1.0 [29]. The result primarily favors genetic-code matches (Figure 4), and secondarily favours single `a↔g` or `c↔t` mismatches, e.g. asparagine scores +5 with `agc` and −14 with `tca`. The gap scores are *a_I_, b_I_, f_I_, g_I_* = −28, −1, +3, 0 and *a_D_, b_D_, f_D_, g_D_* = −23, −1, +3, 0. So frameshifts are not disfavored, perhaps because RepeatMasker’s proteins are close to the fossils’s most recent active ancestors. The positive *f_I,D_* values might be caused by the gap-length distribution not fitting the simple affine model, with an excess of length-1 and length-4 gaps [33].

**Fig. 4.**
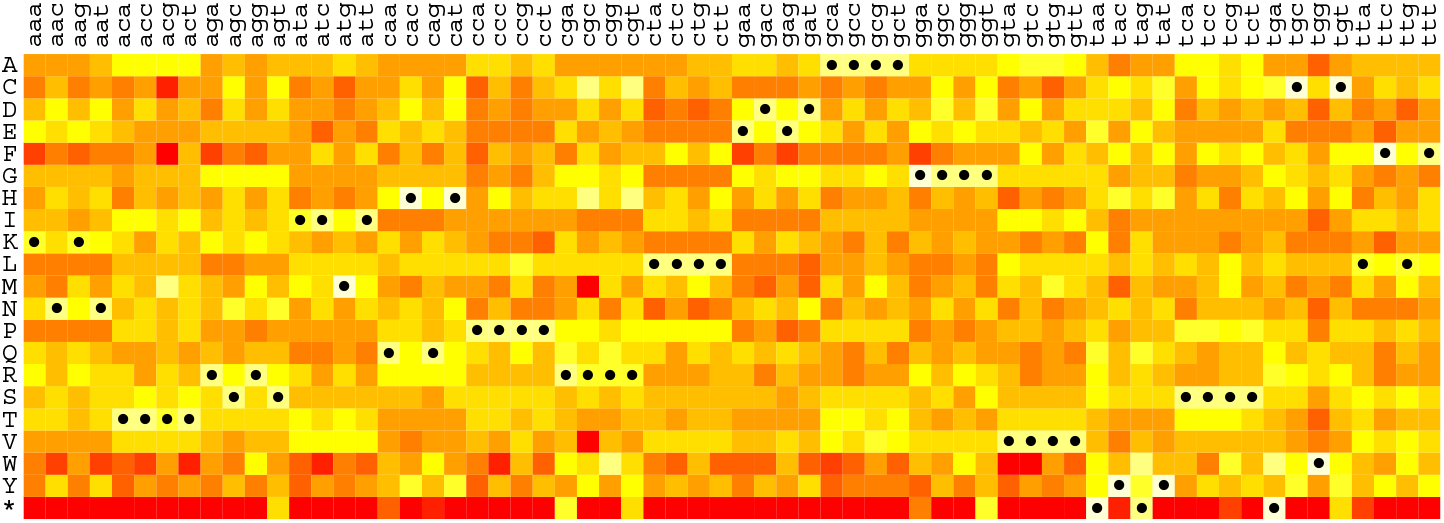
Substitution matrix inferred from human chromosome 21 versus RepeatMasker proteins. Darker red means more disfavored and paler yellow means more favored. Black dots indicate the standard genetic code.

### 3.2 Significance calculation & simple sequences

To test the accuracy of our significance estimates, for the chr21-TE parameters, we calculated *s*_max_ by our full Forward-Backward algorithm for 1000 pairs of random i.i.d. protein and codon sequences, with *Φ_x_* = *ϕ_x_*, *ψ_Y_* = *ψ_Y_*, *M* = 200, and *N* = 602. The observed distribution of *s*_max_ agrees reasonably well with that predicted by Equation 2 (Fig. 5A).

**Fig. 5.**
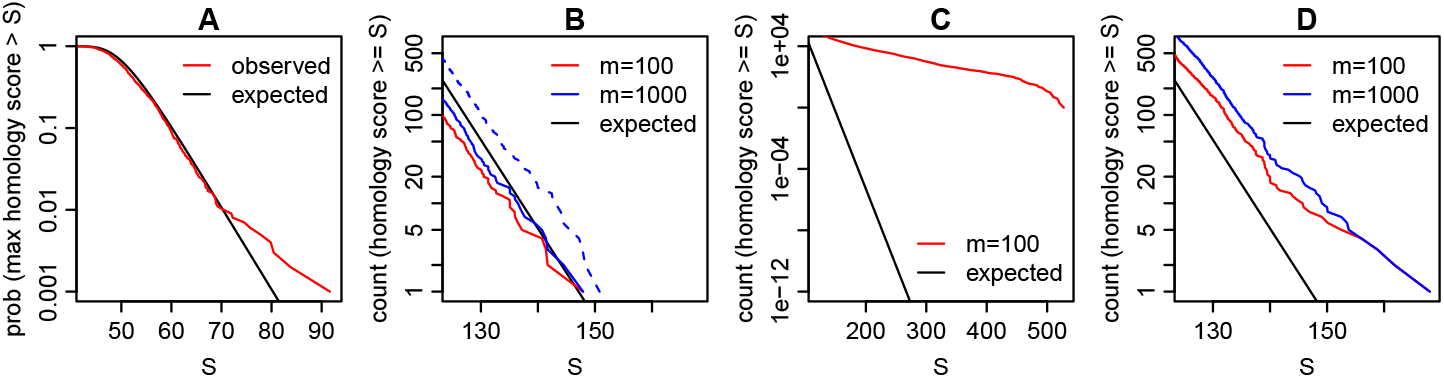
Distributions of homology scores using the chr21-TE parameters. (**A**) Exact homology scores for random protein & codon sequences. (**B**) Seed-and-extend homology scores for random protein & DNA sequences. The dashed line shows a test with (*Φ, ψ*) ≠ (*ϕ, ψ*). (**C**) Seed-and-extend homology scores for TE proteins & reversed chr21 without and (**D**) with simple-sequence masking.

To test whether our significance estimates apply to our seed-and-extend homology search, we compared one pair of random i.i.d. protein and DNA sequences, with *Φ_x_* = *ϕ_x_*, *ψ_y_* = *ψ_y_*, and lengths equal to the number of unambiguous letters in the TE proteins and chr21. The search sensitivity depends on the seed parameter *m*: as *m* increases, sensitivity increases, and the distribution of homology scores approaches the Gumbel prediction (Fig. 5B).

We then considered (*Φ, ψ*) ≠ (*ϕ, ψ*), because the marginal frequencies of *π_xY_* differ from the letter abundances in the TE proteins and chr21, e.g. *ψ*_a_:*ψ*_c_:*ψ*_g_:*ψ*_t_ = 40:19:18:23 but chr21 is 29:21:21:29. So we compared another pair of random i.i.d. protein and DNA sequences, with *Φ_x_* and *ψ_y_* equal to the frequencies in the TE proteins and chr21. In this test, the E-values (expected counts) were too low by a factor of about 3 (Fig. 5B).

Homology search is confounded by “simple sequences”, e.g. ttttcttttttcctt, which evolve frequently and independently. There are various methods to suppress such false homologies, but most do not fully succeed [9, 8]. To illustrate, we compared reversed (but not complemented) chr21 to the TE proteins: this test has no true homologies, but we found many highly-significant homology scores (Fig. 5C). Our solution is to mask the DNA and protein with tantan [9], which eliminates extremely-significant false homologies, at least in this test (Fig. 5D). Further testing is warranted, e.g. here we used a default `tantan` parameter *r* = 0.005, but 0.02 was suggested for DNA-protein comparison [9].

### 3.3 Comparison to blastx

To test whether our homology search is more sensitive than standard methods, we compared chr21 to the TE proteins with NCBI BLAST 2.11.0:

~~~
makeblastdb -in RepeatPeps.lib -dbtype prot -out DB
blastx -query chr21.fa -db DB -evalue 0.1 -outfmt 7 > out
~~~

We repeated this comparison with our method (in LAST version 1177):

~~~
lastdb -q -c -R01 myDB RepeatPeps.lib
last-train --codon -X1 myDB chr21.fa > train.out
lastal -p train.out -D1e9 -m100 -K1 myDB chr21.fa > out
~~~

Option `-q` appends * to each protein; `-R01` lowercases simple sequence with `tantan`; `-c` requests masking of lowercase; `-X1` treats matches to unknown residues (which are frequent in these proteins) as neutral instead of disfavored; `-D1e9` sets the significance threshold to 1 random hit per 10^9^ basepairs; `-m100` sets *m* = 100; `-K1` omits alignments whose DNA range lies in that of a higher-scoring alignment.

This test indicated that our method has much better sensitivity and speed. The single-threaded runtimes were 193 min for `blastx` and 18 min for `lastal`. `blastx` found alignments at 2604 non-overlapping sites on the two strands of chr21, of which all but 23 overlapped LAST alignments. LAST found alignments at 6640 non-overlapping sites, of which 4499 did not overlap `blastx` alignments. All but 21 of LAST’s sites overlapped same-strand annotations by RepeatMasker open-4.0.6 - Dfam 2.0 (excluding Simple_repeat and Low_complexity) [29, 32], suggesting they are not spurious.

### 3.4 Discovery of missing TE orders in the human genome

Eukaryotic TEs have immense diversity, but can be classified into ∼7 major orders: LTR, LINE, and tyrosine-recombinase (YR) retrotransposons, and DDE transposons, cryptons, helitrons, and polintons [36]. Three of them (YR retrotransposons, cryptons, polintons) have not been found in mammals [24, 3].

By comparing the whole human genome (hg38) to RepeatMasker’s TE proteins, we found two of these missing TE orders: YR retrotransposons and polintons. We found polinton alignments at 18 non-overlapping genome sites, with E-values as low as 1.8e-36. Five of these sites are clustered in chromosome 7, indicating that an ancient polinton was fragmented by insertion of an LTR element and an inversion (Fig. 6). We found both major superfamilies of YR retrotransposon: DIRS at 20 non-overlapping sites with min E-value 3e-45, and Ngaro at 4 non-overlapping sites with min E-value 5.1e-14.

**Fig. 6.**
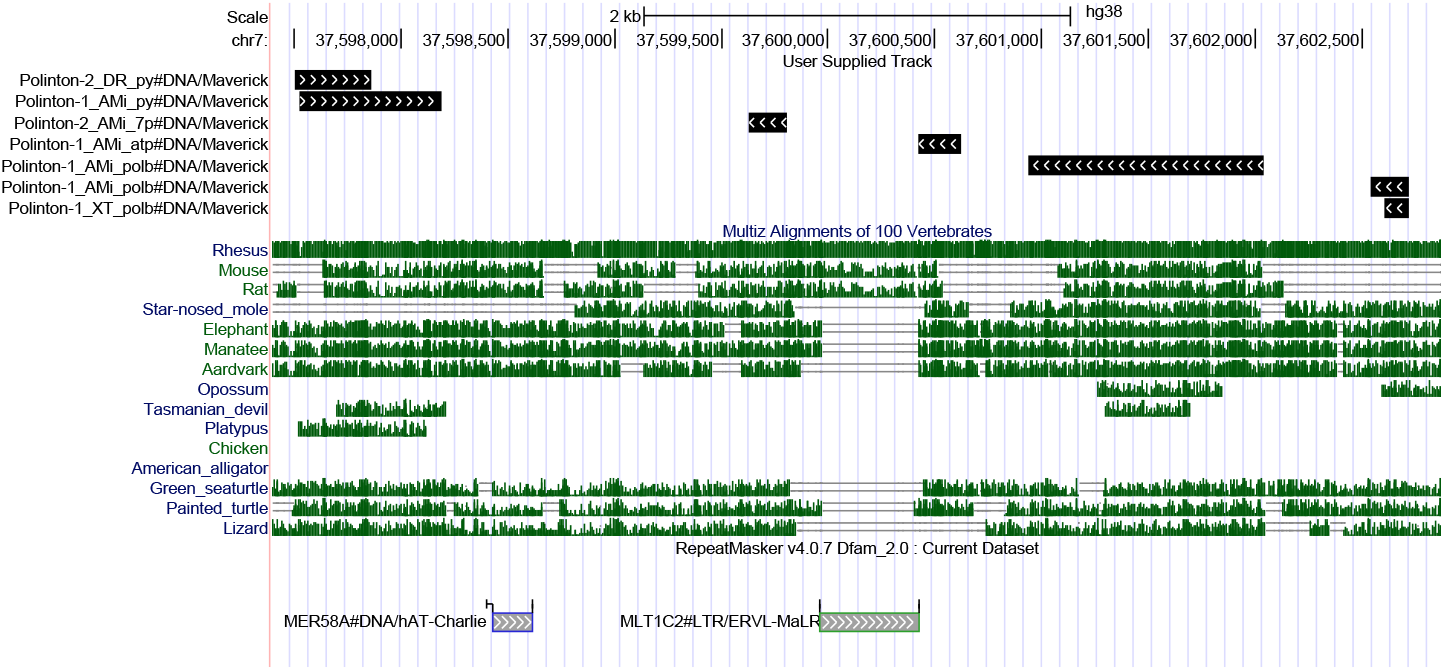
An ancient polinton in human chromosome 7. Black bars: alignments to polinton proteins, arrows indicate +/− strand. Green: alignments to other vertebrate genomes [14]. Screen shot from http://genome.ucsc.edu [18].

## 4 Discussion

Our DNA-protein homology search method seems to be fast, specific, and highly sensitive. It should enable discovery of more ancient and subtle fossils, such as the human polinton, DIRS and Ngaro elements found here. So almost all known major TE categories have left traces in the human genome, suggesting an ability to spread broadly among eukaryotes.

Possible future improvements include better seeding, and using position-specific information on variability of a sequence family [5, 38]. Our significance calculation becomes inaccurate for short sequences, so a finite size correction would be useful [37]. Our method’s parameter-fitting makes it versatile, but it would be better to use different parameters for fossils of different ages.

The sum-of-paths and significance paradigm is not specific to DNA-protein comparison, so could benefit homology search generally. A previous study made similar conjectures on significance of probabilistic homology scores [7]. We suspect those conjectures may be too broad: e.g. one set of substitution and gap scores corresponds to a range of probability models with different values of *t* [10], but only one *t* can appear in the Gumbel formula (Equation 2).

## Acknowledgments

We thank the Frith and Asai lab members for discussions that clarified our thinking.

